# An Investigation of the Effects of Nitrate vs. Ammonium on Plants Using Metabolic Modeling

**DOI:** 10.1101/2022.10.11.511848

**Authors:** Iris Lai, C. Y. Maurice Cheung

## Abstract

As metabolism is a very complex process, it is essential to use computational modeling to better understand the changes in metabolism at a systems level. In this paper, we used a computational model that simulates the metabolism of mature plants to test how the use of nitrate and ammonium as nitrogen sources affects plant metabolism. Our model predicted and showed the possible changes in metabolism in different plant tissues in response to the use of different nitrogen sources. Our methodology produces predictions of metabolic fluxes in plants and improves our understanding of how plants react to changes in conditions. This research can potentially give insights into how we can improve crop yield by identifying metabolic processes that are important in plant growth and in the adaptation to different nitrogen sources. In future research work, we can apply the existing model to explore the effects of different nutrient availability under different conditions (e.g. well-watered vs drought) to optimize nutrient availability under particular environmental conditions. Experimental researchers, plant breeders, and farmers can use the knowledge gained from the modeling work to further our understanding of plant metabolism and apply the knowledge to improve plant growth in the lab and the field.

## Introduction

All plants find their essential nutrients (the three most important being phosphorus, nitrogen, and potassium) in the soil. Nitrogen, arguably the most vital nutrient, is an important component of the amino acids that form the building blocks of plant proteins and enzymes. Proteins and enzymes are a key part of a plant’s cellular machinery, which refers to both the physical and chemical factors of a cell that functions together to ensure that the physiological functions of the cell are carried out. Nitrogen is also an essential nutrient in nucleic acids, which create the cell’s genetic material. The genetic material of plant cells, like all other living organisms, is deoxyribonucleic acid, better known as DNA (Grotewold et al. 2015). DNA determines the features of the plant, including size, color, shape, and even pest resistance, hence the creation of genetically modified plants. Nitrogen can be sourced in various forms. This nutrient can be found in nitrogen fertilizer, which is widely used by farmers and can contain one or more of four sources of nitrogen: nitrate, ammonia, ammonium, or urea (Mengel, 1986). However, the two main sources of nitrogen for plants are nitrate (NO_3_^-^) and ammonium (NH_4_^+^). Most plants prefer NO_3_^-^ as their main source of nitrogen, but a few prefer NH_4_^+^. Though ammonium is an important part of many metabolic processes in plants, many plants experience symptoms of ammonium toxicity when encountering high amounts of NH_4_^+^ such as leaf chlorosis, root hair disfigurement, seedling death, and root axis shrinkage (Pan et al., 2016). Nevertheless, nitrate and ammonium are both inorganic nitrogen sources that are taken up by plant roots to create the amino acids glutamine and glutamate, which are then used to synthesize other amino acids. In order to produce glutamine, nitrate and ammonium need to be converted to ammonia, which combines with glutamate to create glutamine (Taiz and Zaiger, 2010). Even though nitrate is the more popular source of nitrogen for plants, it takes far more energy to convert nitrate to ammonia (nitrogen fixation) compared to ammonium to ammonia. Traces of both nitrogen sources can be found where plants reside, but varying levels of each exist in different locations.

Metabolic modeling can be used to investigate the impacts of varying levels of nitrate and ammonium on plant metabolism. Metabolic modeling is the use of mathematics to represent the metabolism of an organism (Wang et. al., 2019). This innovative way of studying plant metabolism was made possible by the development of modern-day computational power and the vast amount of accumulated information about enzymes, which is a key component of metabolic systems. With metabolic modeling, we can study metabolism at a systems level by studying the behavior of the entire metabolic system instead of studying single reactions. The metabolic system consists of hundreds or even thousands of reactions and metabolites. Metabolic reactions, such as those involved in breaking down or synthesizing biomolecules, result in key metabolites and biomolecules that keep a metabolic system running. Metabolites are the biomolecules involved in the numerous reactions that occur in metabolism, such as lipids, amino acids, and sugars. They can also help metabolism by contributing to energy conversion, signaling, and epigenetic influence (Wellen and Thompson, 2012). Metabolic modeling is an efficient way to predict the outcomes of plant metabolism and its impacts when factors such as nitrogen sources and water levels are altered.

There are little to no other studies using metabolic modeling that show the effect of assorted levels of the two main nitrogen sources, nitrate and ammonium. Thus, to study the complex metabolic system in plants, we use metabolic modeling to test and understand how varying levels of nitrate and ammonium affect the behavior of the metabolic system in plants. Using the model, we can predict the outcome of the impact of the different nitrogen source levels and collect the data to understand its influence on the plants better. It is possible that the outcomes of this research can give more insight into producing better crop yields by identifying important processes in metabolism affected by different nitrogen conditions. With this information, farmers can be better educated on the optimal nitrogen sources and change their techniques for crop growth to optimize their yield.

## Methodology

To run the model simulations and get predictions of metabolic fluxes from metabolic models, a modeling method called flux balance analysis (FBA) was used. FBA is a mathematical approach to predicting the metabolic fluxes (flow of metabolites) in a metabolic system (Orth et. al., 2010). This approach to modeling requires the application of constraints and an objective function. Constraints set the minimum and maximum range of values of the metabolic fluxes for each reaction. In this study, the constraints used were similar to those in a similar study on modeling plant metabolism (Shaw and Cheung, 2018). Sucrose and amino acids were constrained to be exported from the mature leaf. The sucrose and amino acid transport during the light to dark phases were constrained at a ratio of 3:1, i.e. the output of the model is constrained to occur three times faster in the day compared to the night. Cellular maintenance costs were taken into account in the constraints with the ATP and NADPH maintenance costs ratio of 3:1 as in Shaw and Cheung, (2018). An objective function was also needed in FBA to obtain biologically relevant predictions of metabolic fluxes. Our model’s primary objective function was set to minimize photon requirements to produce sucrose and the amino acids to the phloem. A python package, scobra (https://github.com/mauriceccy/scobra), an extension of COBRAPy (Ebrahim et. al., 2013) - a popular python package for FBA, was used to run the simulations of the metabolic model in this study. All scripts in the study were written in Python (www.python.com) and are available in Supplementary Data 1.

The core plant metabolic model that was used in this study is a stoichiometry model. In other words, the model contains information on the stoichiometry of the reactions. In total, there are 1,349 reactions that are contained in this model including reactions representing the light and dark phases (Supplementary Data 2). The core plant metabolic model was used to model the metabolism of mature leaves of C_3_ and CAM plants (Cheung and Shaw, 2018). In this study, we used the model to investigate the use of various levels of nitrate and ammonium as nitrogen sources and made predictions on the resulting metabolic fluxes.

## Results and Discussions

### Metabolic modeling showed changes in metabolism with different nitrogen sources

The metabolic model used included core reactions that simulated the photosynthetic system in plants and can be used to predict the metabolic fluxes when different factors in the environment were changed. The model is based on *Arabidopsis thaliana*, the most well-studied model plant, but it includes core reactions that are present in C_3_, C_4_, and CAM plants. This model simulates a mature plant producing sucrose and amino acids which are transported to the phloem. Thus, the model’s output was constrained to producing sucrose as well as numerous amino acids that would be exported to the phloem and was based on the typical amount of those components in C_3_ plants (Tay et al., 2021). In this scenario, the levels of nitrate and ammonium were adjusted to different levels to find which reactions and metabolites were impacted when either one of the nitrogen sources was at higher levels. Using flux-balance analysis, the metabolic fluxes from using only nitrate or only ammonium as the nitrogen source were predicted. In total, 101 reaction fluxes changed out of 1349 reactions with 46 being metabolic reactions and the rest being transport reactions. The full flux solutions of the two scenarios can be found in Supplementary Data 3.

Various interesting changes in metabolic fluxes were observed. Selected reactions with flux changes between the two scenarios can be found in Table 1. From the changes in fluxes of the external exchange reactions, we can see that there is an increase in photon demand in the nitrate-only scenario compared to the ammonium-only scenario. This can be explained by the extra energy and reducing power needed to reduce nitrate to ammonium before the incorporation of ammonium into amino-acids. As expected, reactions involved in nitrate reduction were only active in the nitrate-only scenario. This leads to some system-wide changes in energy and reductant metabolism. It was predicted that the photosynthetic electron transport chain carried more flux in the nitrate-only scenario given the increase in energy and reductant demand. This also impacted the shuttling of reductant across subcellular compartments with less reductant shuttled out of the chloroplast in the nitrate-only scenario as can be seen by the fluxes of malate dehydrogenases (Table 1). The knock-on effect is that less reductant is available for the mitochondrial electron transport chain which has a lower flux in the nitrate-only scenario compared to the ammonium-only scenario. Interestingly, with the shift on reductant demand and shuttling, which also affect ATP synthesis, the model predicted a shift in glutamine synthetase flux with more glutamine synthetase flux going through the chloroplast in the nitrate-only scenario. One possible explanation is that with the increase in reductant demand by using nitrate as the nitrogen source, to maintain the NADPH to ATP demand ratio in the chloroplast, more ATP is used by glutamine synthetase in the chloroplast in the nitrate-only scenario. This example is a good demonstration of how large-scale metabolic models can give insight into the systems-level behavior of the metabolic system.

**Table 1.** Selected key reactions with changes in flux values between ammonium-only and nitrate-only scenarios

| Reaction types | Reaction name in model | Ammonium only | Nitrate only |
| --- | --- | --- | --- |
| External exchange reactions | Photon_tx1 | 178.7729851 | 180.0416573 |
|  | Nitrate_tx1 | 0 | 0.113274306 |
|  | NH4_tx1 | 0.113274306 | 0 |
|  | H_tx1 | 0.225952236 | -0.000596376 |
|  | H2O_tx1 | 11.53141222 | 11.64468652 |
|  | O2_tx1 | -10.41376376 | -10.64031237 |
| Nitrate reduction | FERREDOXIN_NITRITE_REDUCTASE_RXN_p1 | 0 | 0.113274306 |
|  | NITRATE_REDUCTASE_NADH_RXN_c1 | 0 | 0.113274306 |
| Photosynthetic electron transport | RXN490_3650_p1 | 89.38649254 | 90.02082865 |
|  | PLASTOQUINOL_PLASTOCYANIN_REDUCTASE_RXN_p1 | 44.69324627 | 45.01041433 |
|  | 1_PERIOD_18_PERIOD_1_PERIOD_2_RXN_p1 | 42.72685244 | 42.70419758 |
|  | PSII_RXN_p1 | 22.34662314 | 22.50520716 |
|  | Plastidial_ATP_Synthase_p1 | 19.1542484 | 19.29017757 |
| Malate-OAA reductant shuttles | MALATE_DEHYDROGENASE_NADPs_RXN_p1 | 5.827002303 | 5.508277734 |
|  | MALATE_DEH_RXN_m1 | 2.450807036 | 2.314877869 |
|  | MALATE_DEH_RXN_c1 | 1.454846665 | 1.272051263 |
| Mitochondrial electron transport | NADH_DEHYDROG_A_RXN_mi1 | 4.326852938 | 4.190923771 |
|  | Mitochondrial_ATP_Synthase_m1 | 4.514457528 | 4.378528361 |
| Glutamine Synthetase | GLUTAMINESYN_RXN_p1 | 0.001802277 | 0.113520071 |
|  | GLUTAMINESYN_RXN_m1 | 1.987763695 | 1.876045902 |

Another key change in flux is the external exchange of protons with the ammonium-only scenario having a significant export of protons while the nitrate-only scenario has close to zero flux in proton exchange with the environment. This can have a significant impact on the pH balance of the system as using ammonium requires the plant to export the excess protons to the environment, which can be energetically costly and could also impact the pH of the environment, e.g. soil around the plant.

### Varying amount of nitrate and ammonium showed linear change in metabolism

We conducted a scan by varying the amount of nitrate as the nitrogen source and predicted the resulting metabolic fluxes with the change in nitrate influx. One of the interesting metabolic changes seen is the increasing amount of photons as the amount of nitrate increases (Figure 1), which is consistent with our results earlier. We can also see the increase in plastidial ATP synthase flux and the decrease in mitochondrial ATP synthase flux as nitrate influx increases (Figure 1). Consistent with our earlier results, there is a decrease in proton export as more nitrate is used (Figure 1). More importantly, it can be seen from the results of the scan that the changes in flux, especially for the ATP synthases, occur linearly with an increase in nitrate influx, suggesting a lack of complex metabolic rewiring that impacts energy metabolism. Our results suggest that plants can readily switch from using ammonium to nitrate without complex rewiring of metabolism, but the change in nitrogen sources do have an impact on energy and reductant metabolism as well as proton export, which could have potential impact on the pH balance of the system.

**Figure 1.**
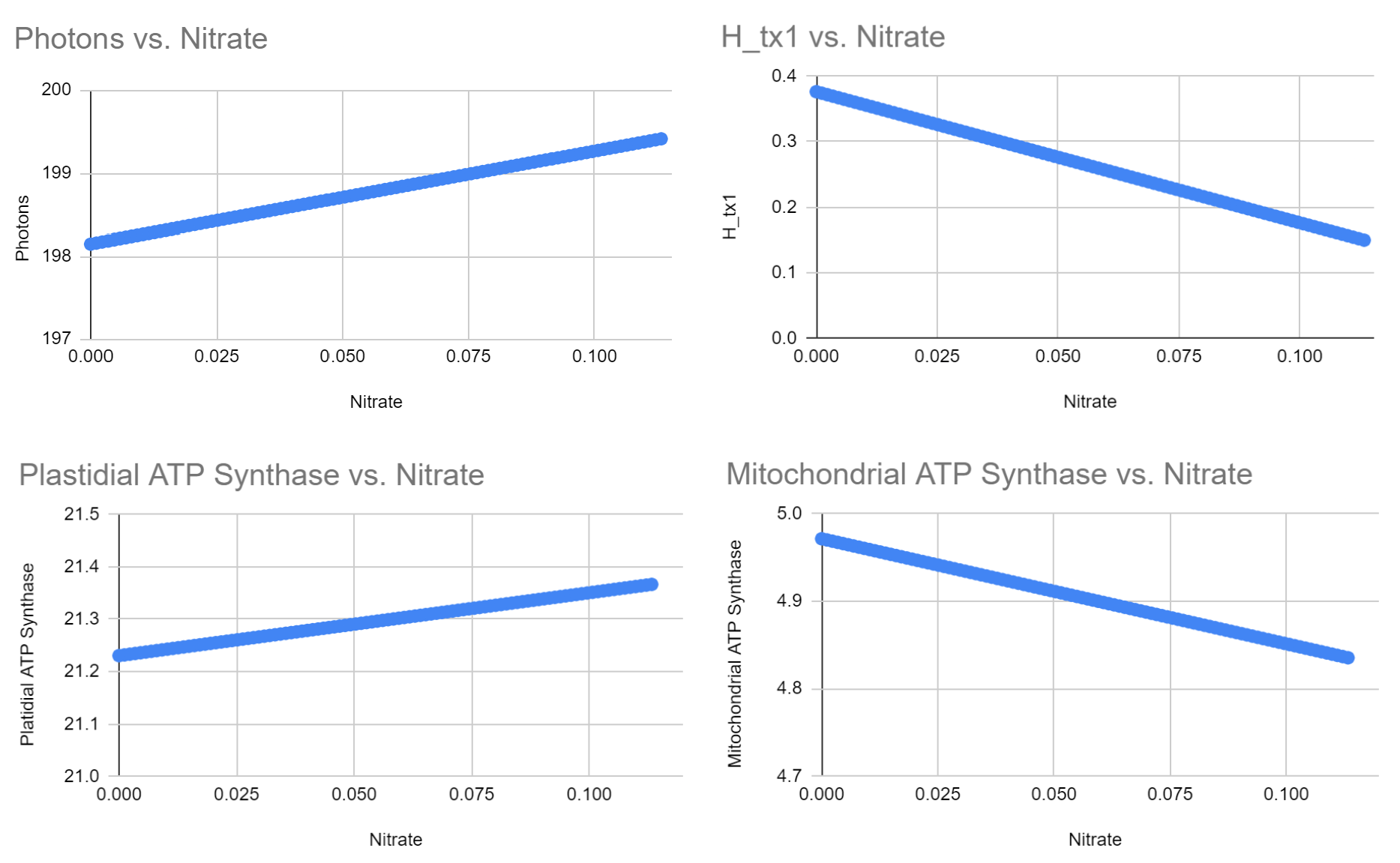
Fluxes of photon influx, external proton export, plastidial ATP synthase, and mitochondrial ATP synthase as nitrate influx increases.

## Conclusion

Using metabolic modeling, it is possible to study the metabolic changes resulting from the variability in resource availability. In this study, two variations of nitrogen source levels were investigated, and predictions were made of the change in reactions based on whether a mature plant received nitrogen from only nitrate or only ammonium. With this knowledge, farmers and metabolic engineers can improve nitrogen efficiency.

## Supplementary Material

Supplementary Data 1 - Python scripts used for simulations

Supplementary Data 2 - Flux balance core plant metabolic model

Supplementary Data 3 - Flux solutions for ammonium-only and nitrate-only scenarios

Supplementary Data 4 - Flux solution from nitrate-ammonium scan

